# EPEK: creation and analysis of an Ectopic Pregnancy Expression Knowledgebase

**DOI:** 10.1101/2022.12.20.521279

**Authors:** Ananya Natarajan, Nikhil Chivukula, Gokul Balaji Dhanakoti, Ajaya Kumar Sahoo, Janani Ravichandran, Areejit Samal

**Affiliations:** Indian Institute of Science Education and Research (IISER) Mohali, Punjab, India; The Institute of Mathematical Sciences (IMSc), Chennai, India; Homi Bhabha National Institute (HBNI), Mumbai, India

**Keywords:** Ectopic Pregnancy, expression profile, EPEK resource, enrichment analysis, network analysis

## Abstract

Ectopic pregnancy (EP) is one of the leading causes of maternal mortality, where the fertilized embryo grows outside of the uterus. Recent experiments on mice have uncovered the importance of genetic factors in the transport of embryos inside the uterus. In the past, efforts have been made to identify possible gene or protein markers in EP in humans through multiple expression studies. Although there exist comprehensive gene resources for other maternal health disorders, there is no specific resource that compiles the genes associated with EP from such expression studies. Here, we address that knowledge gap by creating a computational resource, Ectopic Pregnancy Expression Knowledgebase (EPEK), that involves manual compilation and curation of expression profiles of EP in humans from published articles. In EPEK, we compiled information on 314 differentially expressed genes, 17 metabolites, and 3 SNPs associated with EP. Computational analyses on the gene set from EPEK showed the implication of cellular signaling processes in EP. We also identified possible exosome markers that could be clinically relevant in the diagnosis of EP. In a nutshell, EPEK is the first and only dedicated resource on the expression profile of EP in humans. EPEK is accessible at https://cb.imsc.res.in/epek.

## 1. Introduction

Ectopic Pregnancy (EP) is a pregnancy complication where the fertilized embryo implants on a region other than the uterus. EP occurs in almost 2% of all global pregnancies [1,2] and accounts for 10-15% of all maternal deaths during the first trimester [3,4]. The most common extrauterine implantation site is the fallopian tube, accounting for more than 98% of EP cases [1,2]. Early diagnosis of EP is necessary for preventing adverse health outcomes resulting from tubal rupture, such as pelvic pain, vaginal bleeding and ultimately death [5]. Initial diagnostic treatments for EP include prescription of methotrexate to stop the embryo from growing at the implantation site whereas in the case of ruptured EP a surgical intervention becomes inevitable [6,7]. Nonetheless, the early diagnosis of EP is challenging as the clinical symptoms do not manifest until the tubal damage [5]. Therefore, it is imperative to understand the pathophysiology of EP in order to identify novel therapeutic targets.

Several risk factors such as smoking, history of EP, tubal damage, use of intrauterine device, pregnancy conceived by assisted reproduction, pelvic inflammatory disease, and a history of abortions are reported to contribute to the pathogenesis of EP [6,8]. It is speculated that these risk factors aid in the dysfunction of the embryo transport mechanism or of the general physiology of the tubal environment [2]. In a recent experimental study on mice, Bianchi *et al*. [9] have provided novel insights into the molecular mechanisms involved in the maintenance of tubal embryo transport. In contrast, such comprehensive studies cannot be carried out on humans due to ethical constraints. In this scenario, comparative studies give a descriptive understanding of the differences in the expression profiles of ectopic and nonectopic conditions.

Knowledgebases built on comparative studies are helpful in deciphering the underlying pathophysiology of various human diseases. In particular, different knowledgebases built for maternal health disorders relied on the differential expression of genes or proteins [10–13]. Such data-driven knowledgebases have highlighted the molecular mechanisms implicated in these diseases. Though several comparative studies on the differential expressions in EP have been carried out [14–17], there is no unique computational resource that collates all the experimental information on expression profiles of EP from the published literature.

Therefore, to fill the knowledge gap, we built Ectopic Pregnancy Expression Knowledgebase (EPEK), a manually curated resource on expression profiles of EP. EPEK compiles information on the expression profile of EP from published articles with experimental evidence. In addition, EPEK also provides curated information on metabolite profiles and single nucleotide polymorphisms (SNPs) associated with EP. To the best of our knowledge, EPEK is the first dedicated resource on ectopic pregnancy. EPEK is available for academic research at: https://cb.imsc.res.in/epek.

Additionally, we employed several computational approaches to analyze the gene set compiled in EPEK. Initially, we performed different enrichment analyses to understand the different biological functions, processes and risk factors associated with EP. Further, a protein-protein interaction (PPI) network revealed the probable molecular interactions in the gene set associated with EP. Finally, a comparison with other maternal health disorders highlighted the novelty and uniqueness of EPEK.

## 2. Methods

### 2.1. Workflow to compile and curate relevant articles associated with ectopic pregnancy

Our primary objective is to compile the expression profiles of genes, proteins, and miRNAs associated with EP from published research articles. In addition, we also compile the SNPs and metabolite profiles associated with EP from published research articles. To search relevant published articles for EP, we designed a query to mine PubMed (https://pubmed.ncbi.nlm.nih.gov/) that also incorporates the different Medical Subject Headings (MeSH) terms (https://www.ncbi.nlm.nih.gov/mesh/) associated with EP, namely, Extrauterine Pregnancies, Tubal Pregnancies, and Abdominal Pregnancies. We performed the PubMed search using the following query:

‘((ectopic OR extrauterine OR tubal OR abdominal) AND (pregnancy OR pregnancies)) AND ((gene* OR genom* OR microarray OR transcriptom* OR (gene* AND expression)) OR (protein* OR proteom* OR (gene AND product*) OR (protein* AND expression)) OR (metabolite* OR metabolom* OR metabolic OR metabolize OR metabolise) OR ((“single nucleotide” AND (polymorphism* OR variation*)) OR SNP* OR (gene* AND (variation* OR mutation* OR polymorphism*))) OR (miRNA OR microRNA))’.

This PubMed search was last done on 22 December 2021 and led to the retrieval of 2857 published research articles (Figure 1). Thereafter, we manually screened the 2857 articles to curate the list of relevant studies on EP (Figure 1).

**Figure 1:**
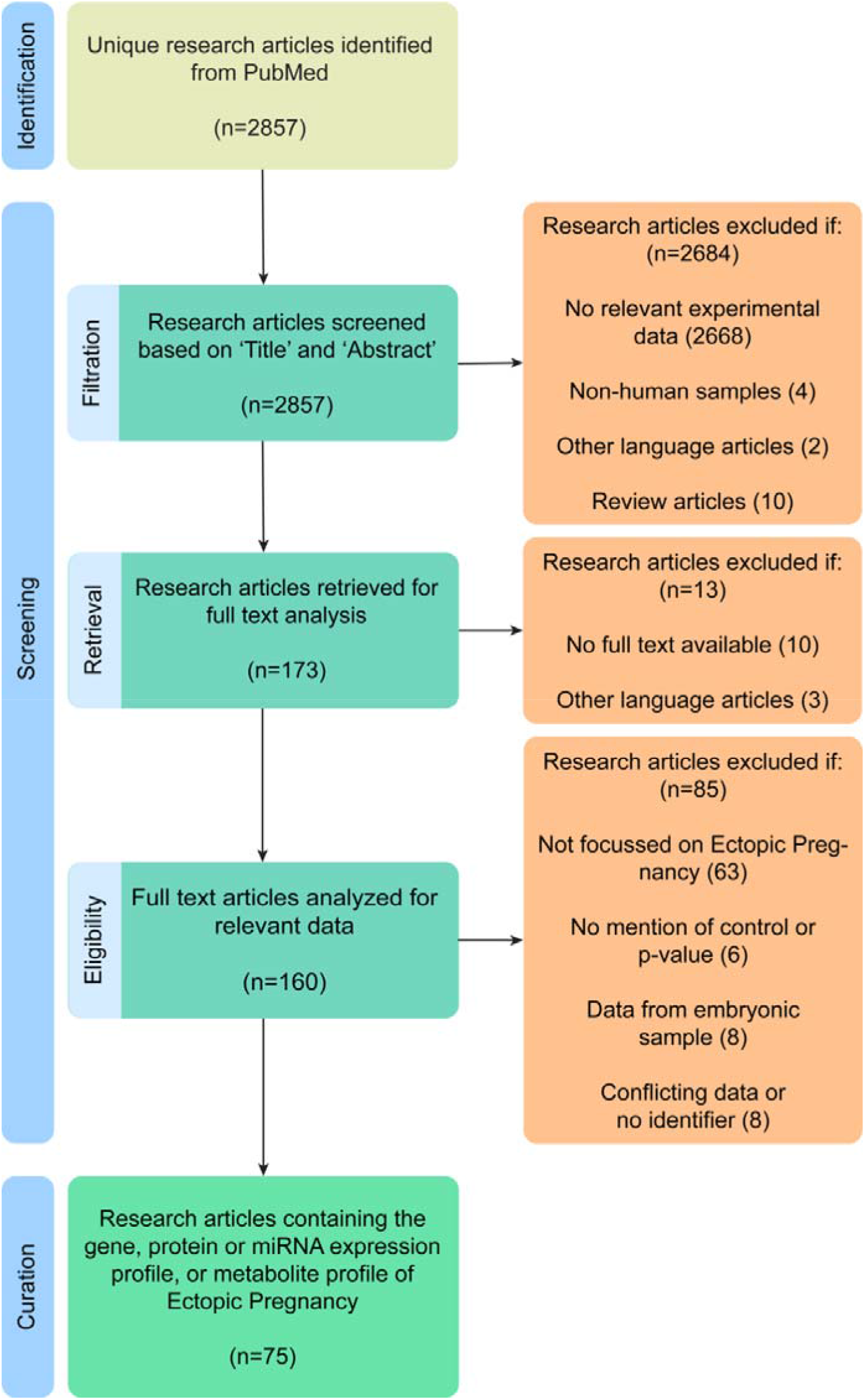
Workflow to identify published research articles containing information on expression profiles of gene, protein and miRNA, metabolite profiles and SNPs associated with ectopic pregnancy (EP) in humans. This workflow is presented according to PRISMA statement [18].

Initially, we screened the 2857 articles by considering only the title and abstract. In this step (Figure 1), we excluded an article if: (i) the article does not seem to contain any experimental data relevant to EP; (ii) the article does not report experiments performed on human samples; (iii) the article is written in language other than English; (iv) it is a review article. At the end of this initial screening step, we identified 173 published articles, and thereafter, we proceeded to collect the full text of these articles. In this step, we excluded 13 more articles as their full text was unavailable or written in a language other than English (Figure 1). Subsequently, we read through the full text of the remaining 160 articles and excluded articles if: (i) they are not focused on EP; (ii) there is no mention of control groups in the experiment or experimental results are not reported as significant; (iii) expression data was collected from embryo instead of maternal tissue (Figure 1). Furthermore, we omitted studies if the reported results contradicted each other.

Finally, we curated 75 research articles (Figure 1) which contain relevant data associated with EP. Figure 1 presents the complete workflow to compile and curate the relevant articles associated with EP according to the PRISMA statement [18]. Additionally, we searched the GEO [19] database (https://www.ncbi.nlm.nih.gov/geo/) for gene expression profiles associated with EP, and the two relevant studies identified in GEO were already included in our curated set of 75 published articles. We also searched the ClinVar [20] database for clinically significant SNPs associated with EP, but did not find any relevant records.

### 2.2. Data compilation and annotation

From the curated set of 75 research articles, we compiled the data on gene expression profile, protein expression profile, miRNA expression profile, metabolite profile, and SNPs associated with EP. Additionally, we gathered information on the tissue type, experimental techniques used for the study, expression levels, p-value, fold change, type of non-ectopic control, size of control cohort, and size of samples. Thereafter, we annotated the compiled information associated with EP by mapping them to standard databases.

We manually mapped the gene, protein and miRNA records to their respective gene identifiers in NCBI (https://www.ncbi.nlm.nih.gov/gene/) and ENSEMBL [21]. Further, we gathered the corresponding UniProt [22] identifiers for both gene and protein records, the CAS identifiers from ChemIDplus (https://chem.nlm.nih.gov/chemidplus/) for metabolite records, and the SNP identifiers from dbSNP (https://www.ncbi.nlm.nih.gov/snp/) for SNP records. Notably, synonymous gene, protein and miRNA records were grouped together and mapped to the non-redundant official NCBI gene identifiers. Further, we excluded any compiled information on gene, protein, miRNA or metabolite that could not be mapped to identifiers in any standard databases.

At the end of this tedious step on data compilation, curation and annotation, we obtained 314 unique gene identifiers (Supplementary Table S1), 17 unique metabolite identifiers (Supplementary Table S2), and 3 unique SNP identifiers (Supplementary Table S3) associated with EP. We remark that this compiled set of 314 genes associated with EP was subsequently used as ‘input gene set’ to perform various analyses in this work.

For the input gene set, we obtained the functional class information from the PANTHER [23] database and Gene Ontology (GO) terms using the ‘Generic GO Slim’ version in GOnet [24] web-application (https://tools.dice-database.org/GOnet/). Using the KEGG mapper conversion tool (https://www.kegg.jp/kegg/mapper/convert_id.html), we mapped the NCBI identifiers for the input gene set to their corresponding KEGG identifiers. Thereafter, we gathered the information on KEGG [25,26] pathways for the 314 genes using the KEGG mapper search tool (https://www.kegg.jp/kegg/mapper/search.html). Lastly, we also collected the ReconMap3 [27] and KEGG identifiers for the 17 metabolites associated with EP.

### 2.3. Web interface and database management system

Using the compiled information on EP, we have created an online database with a user-friendly web interface namely, **E**ctopic **P**regnancy **E**xpression **K**nowledgebase (EPEK). EPEK compiles three types of experimental records associated with EP, namely, genes, metabolites, and SNPs. Further, EPEK also gathers annotations for the compiled set of genes, metabolites, and SNPs associated with EP, such as identifiers from different databases, pathway information and GO terms. EPEK is openly accessible for academic research at: https://cb.imsc.res.in/epek/.

In the web interface of EPEK, we used MariaDB (https://mariadb.org/) to store the compiled data and Structured Query Language (SQL) to retrieve from the compiled data. To create the web interface, we used PHP (https://www.php.net/) with custom HTML, CSS, jQuery (https://jquery.com/), and Bootstrap 4 (https://getbootstrap.com/docs/4.0/). EPEK database is hosted on an Apache (https://httpd.apache.org/) web server running on Debian 9.4 Linux Operating System.

### 2.4. GO term enrichment analysis

To comprehend the biological significance of the compiled expression profiles associated with EP, we performed GO term enrichment analysis for the input gene set. To perform the GO term enrichment analysis, we used GOnet [24] web-application to get the significantly enriched (p-value ≤ 0.01) GO terms in the input gene set for the 3 ontologies namely, biological process, molecular function and cellular component. For each of the 3 ontologies, we retrieved the significantly enriched GO terms along with the number of genes associated with EP for each GO term (Supplementary Table S4). We ranked the GO terms based on the number of associated genes, and thereafter, selected the top 10 GO terms for visualization. We visualized the top 10 GO terms for each ontology via a bar plot using seaborn [28,29] package version 0.12.1.

### 2.5. Pathway enrichment analysis

We performed pathway enrichment analysis on the input gene set to determine the enriched biological pathways. For this analysis, we relied on the pathway associations captured in the ‘KEGG 2021 Human’ gene set library of Enrichr [30] (https://maayanlab.cloud/Enrichr/). This enrichment analysis resulted in a ranked list of pathways based on the extent of similarity (quantified by a p-value) between the pathway gene set and the input gene set. Here, we considered KEGG pathways with gene set size > 50 to avoid overestimation of the resultant statistics. Moreover, we considered a p-value cut-off of < 0.01 to shortlist 51 enriched KEGG pathways (Supplementary Table S5) in the input gene set. Further, we manually mapped each of these 51 pathways to their corresponding hierarchical classification (Supplementary Table S5) provided by the KEGG BRITE database (https://www.genome.jp/kegg/brite.html). We visualized the enriched pathways in the input gene set via bubble plots generated using seaborn [28,29] package version 0.12.1. We also generated a pathway similarity network by computing the Jaccard index between pairs of pathways based on their gene sets. In the pathway similarity network, two pathways were connected by an edge if the corresponding Jaccard index for the pair is > 0.1. The pathway similarity network was visualized using Cytoscape version 3.9.1 [31].

### 2.6. Functional enrichment analysis using protein-protein interaction network

To identify the functional interactions between genes from the input gene set, we generated protein-protein interaction (PPI) network using NetworkAnalyst 3.0 [32] webserver (https://www.networkanalyst.ca/NetworkAnalyst/home.xhtml). NetworkAnalyst provides tissue-specific PPI networks using information from DifferentialNet [33] database. The NetworkAnalyst webserver allows users to select the tissue type and retrieve the interactions specific to a tissue. While performing this analysis on NetworkAnalyst webserver, a filter value of 1 results in PPIs specific to the selected tissue whereas a filter value of 30 results in PPIs unchanged across tissues. For our analysis, we selected ‘uterus’ as the tissue type and set the filter value to 1 to generate a uterus-specific PPI network from NetworkAnalyst for the input gene set. Additionally, we used the Genotype-Tissue Expression (GTEx) [34] database to filter the uterus-specific nodes in the PPI network. From this resulting PPI network, we generated a first-order network (wherein at least one node in each PPI pair belongs to the input gene set) and a zero-order network (wherein both nodes in each PPI pair belong to the input gene set). Finally, we visualized the largest connected component of the first-order network in conjunction with the zero-order network (Supplementary Table S6) using Cytoscape [31].

### 2.7. Disease enrichment analysis

Similar to the pathway enrichment analysis, we also performed disease enrichment analysis to determine the enriched diseases in the input gene set. For this analysis, we used the ‘CURATED’ gene-disease associations from the DisGeNET [35] database. Note that DisGeNET compiles these ‘CURATED’ associations from PsyGeNET [36], UniProt [22], OrphaNet [37], CGI [38], CTD (human data) [39], ClinVar [20], and the Genomics England PanelApp [40]. Further, DisGeNET provides Gene-Disease Association (GDA) score and Evidence Index (EI) for different associations in the database. According to DisGeNET, the GDA score accounts for the number of sources, type of sources, and number of published articles supporting a gene-disease association, and the EI indicates the contradictions in the gene-disease association among different articles. For our analysis, we considered a GDA score cut-off of > 0.3 and EI cut-off of > 0.5 to filter high confidence gene-disease associations from DisGeNET that have at least 50% of their supporting articles validating the associations. We performed the Fisher’s exact test [41] using the filtered gene-disease associations from DisGeNET as background data to find the enriched diseases in the input gene set. The p-values resulting from the Fisher’s exact test are corrected for false discovery rate (FDR) using the Benjamini-Hochberg method [42]. Moreover, we only considered diseases with gene set size > 50 and p-value cut-off of < 0.01 as in the preceding section to shortlist 49 enriched diseases (Supplementary Table S7) in the input gene set. Similar to the case of enriched pathways, we visualized the enriched diseases via bubble plots and a disease-disease similarity network.

## 3. Results

### 3.1. EPEK: Ectopic Pregnancy Expression Knowledgebase

Our primary aim is to present a highly curated resource of expression profiles associated with EP. To accomplish this objective, we identified 75 published research articles containing relevant information on EP, and thereafter, manually read the full text of these 75 articles to compile the expression profiles associated with EP (Methods; Figure 1). From the published articles, we compiled information on genes identified through transcriptomic studies, proteomic studies and miRNA expression studies, metabolites, and SNPs associated with EP. Further, we collected several annotations for genes, metabolites and SNPs from different databases (Methods). Using standard identifiers, we curated a non-redundant list of 314 genes, 17 metabolites and 3 SNPs associated with EP. Importantly, of the 314 genes associated with EP, 217 are differentially expressed in transcriptomic studies, 120 in proteomic studies, and 7 in miRNA expression studies on EP (Supplementary Table S1). Notably, 30 of these 314 genes are reported to be differentially expressed in both transcriptomic and proteomic studies on EP (Supplementary Table S1). We used this curated data with experimental evidence from the 75 published articles to build the EPEK database that compiles heterogeneous information on the 314 genes (designated as gene products in the EPEK website), 17 metabolites and 3 SNPs associated with EP. In addition, EPEK also compiles information on the GO terms and KEGG pathways for the 314 genes associated with EP. We remark that EPEK is the first and only dedicated resource on expression profiles specific to ectopic pregnancy to the best of our knowledge. EPEK is accessible online at: https://cb.imsc.res.in/epek/.

Figure 2 shows the screenshots of the different pages on the EPEK website. EPEK is an organized online resource that has a friendly web interface with simple navigation (Figure 2a) enabling easy access to the compiled data for the users. Users can query EPEK for: (i) genes based on their identifiers, (ii) genes based on their expression levels, and (iii) metabolites based on their identifiers, to get the respective information (Figure 2b). For each query, EPEK returns the corresponding results in a tabular format (Figure 2c). From the tabulated results, the detailed information page is accessed by selecting the gene symbol or metabolite name (Figure 2d). In addition to the SEARCH option, users can navigate through EPEK using an interactive BROWSE option (Figure 2e). Using the BROWSE option, users can browse for genes, metabolites and SNPs associated with EP, and GO terms and KEGG pathways for the genes in tabular format (Figure 2f).

**Figure 2:**
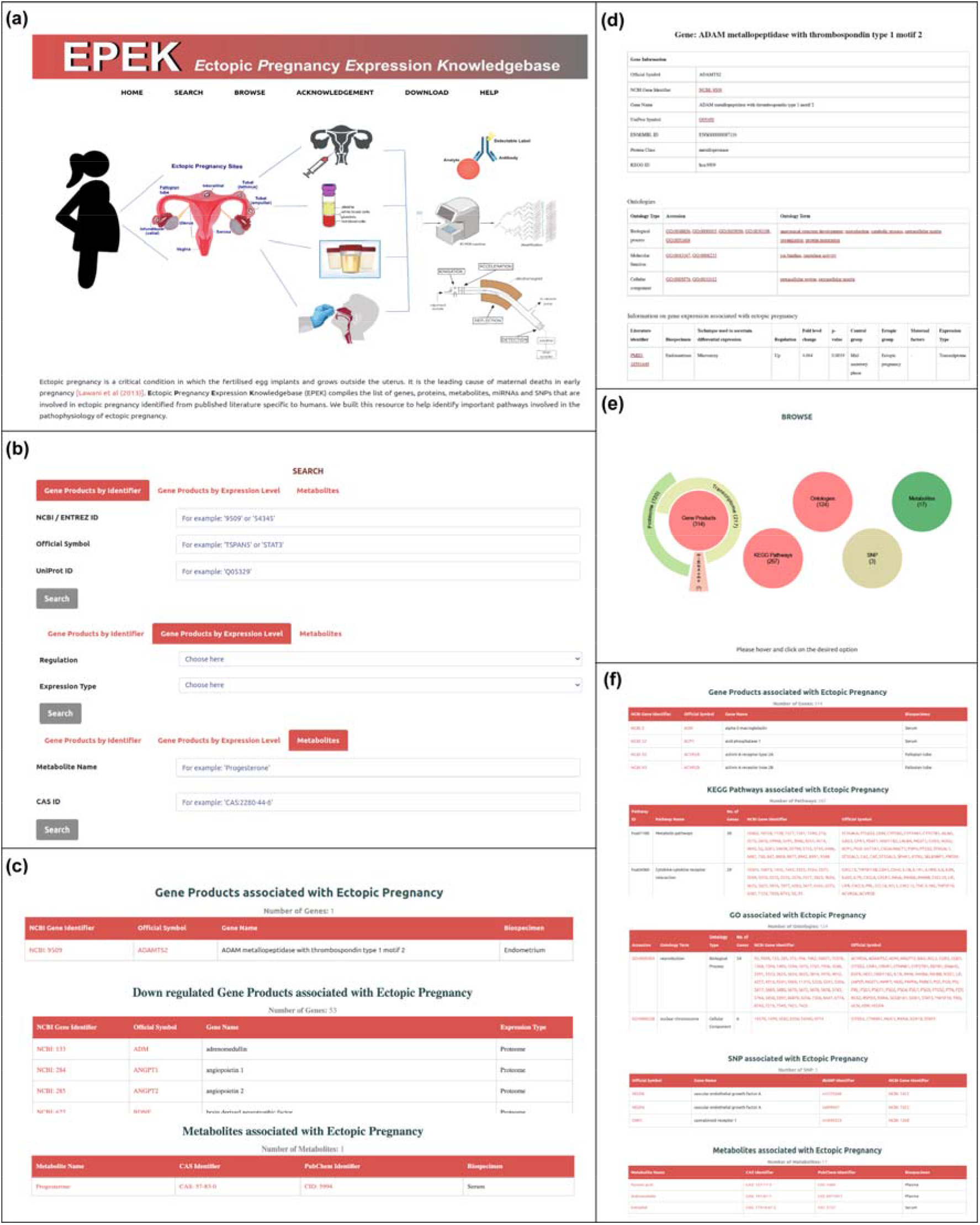
The web interface of EPEK. **(a)** The screenshot of the EPEK homepage with simple navigation bar. EPEK enables users to search the genes and metabolites associated with ectopic pregnancy (EP), browse the genes, metabolites and SNPs associated with EP, and browse GO terms and KEGG pathways for the genes associated with EP by using the SEARCH and BROWSE option. **(b)** The SEARCH option allows users to search genes associated with EP by their names, identifiers and expression levels, and metabolites associated with EP by their names and identifiers. **(c)** Screenshots of the results retrieved in tabular format from the SEARCH option in EPEK. **(d)** Screenshots of the detailed information page for genes or metabolites by selecting the gene symbol or metabolite name in the tabulated results from the SEARCH option in EPEK. **(e)** The interactive BROWSE option enables users to navigate through the genes, metabolites and SNPs associated with EP, and GO terms and KEGG pathways for the genes associated with EP. (**f)** Screenshots of the results retrieved in tabular format by selecting the items in the BROWSE option of EPEK.

### 3.2. Cellular signaling processes are highly enriched in EP

The GO enrichment analysis provides insights into the biological functions associated with a set of genes. In this study, we performed GO enrichment analysis on the 314 genes associated with EP to understand the enriched functions of the genes based on 3 ontologies namely, biological process, molecular function and cellular component (Methods). Figure 3 shows the top 10 enriched GO terms for each of the 3 ontologies. The GO term analysis shows a strong enrichment of cellular signaling processes in the genes associated with EP (Figure 3). In particular, we observed that IL6R gene, which is associated with signaling, is present in most of the GO terms (i.e., 26 out of 30) (Supplementary Table S4). Moreover, previous studies have shown that genes involved in cellular signaling play an important role in the transfer and implantation of eggs in the uterus [2]. In this context, we find a variety of signaling genes (integrins, interleukins, membrane receptors) being differentially expressed in EP.

**Figure 3:**
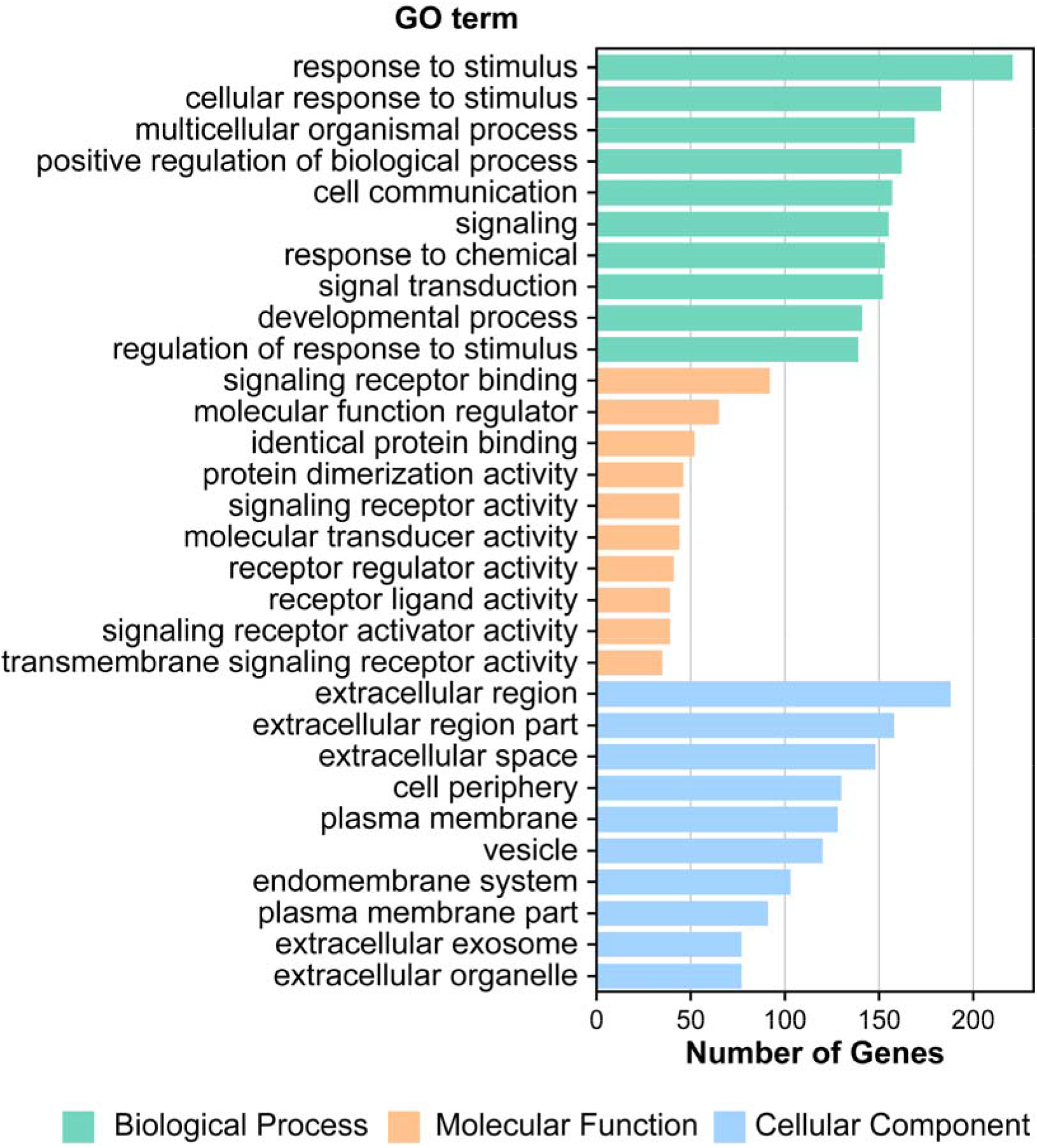
Bar plot of the top 10 enriched (p ≤ 0.01) gene ontology (GO) terms in the 3 ontologies namely biological process (green), molecular function (orange) and cellular component (blue). For each GO term, the horizontal bar gives the number of genes associated with the GO term from the 314 genes associated with EP.

Noting that the GO terms for cellular signaling are significantly enriched in the genes associated with EP, we were further interested to search for clinically relevant biomarkers for EP. Exosomes are extracellular vesicles that act as messengers to carry signals involving a variety of RNAs and proteins [43]. They are released into the extracellular matrix by the cells, and carried in bodily fluids to enable distant communication. The presence of exosomes in the bodily fluids provides for an easy and non-invasive diagnostic study, and therefore invites clinical attention [44]. In this respect, Esfandyari *et al*. [45] have discussed in their review about potential use of exosomes as the source of biomarkers for pregnancy related disorders. Here we explored the presence of any genetic or protein markers present in the exosomes secreted by uterine tissue in our curated list of 314 genes associated with EP. For this, we relied on the uterus and fallopian tubule specific exosome markers from the exoRBase [46] database and the dataset of miRNA markers from the uterine fluid samples compiled by Ng *et al*. [47]. Upon comparison of the 314 genes associated with EP with the two datasets for markers specific to uterus tissue or fallopian tube, we found that 3 of the 7 miRNA namely, MIR323A, MIR324 and MIR517A, and a protein coding gene SNTN are plausible candidates for EP related exosome markers. Further experiments are required to establish the role of the identified miRNAs or protein for exosome related markers in EP.

### 3.3. Immune-response related signaling pathways are highly enriched in EP

Pathway enrichment analysis is a routinely used method to gain mechanistic insights from a set of differentially expressed genes. In this study, we performed pathway enrichment analysis for the 314 genes to determine the pathways that are significantly enriched in the set of genes associated with EP (Methods). Figure 4a shows a bubble plot of the 51 significantly enriched pathways in the input gene set. We observed that the rheumatoid arthritis pathway has the most overlap score and the Cytokine-Cytokine receptor interaction pathway is the most significant pathway (Figure 4a; Supplementary Table S5). Rheumatoid arthritis (RA) is known to be an auto-immune disease where the pathogenesis is attributed to the signaling pathways of different cytokines [48–50]. In the context of RA associated with pregnancy, Clowse *et al*. [51] reported that women with pregnancies after being diagnosed with RA have higher rates of pregnancy related adverse outcomes such as miscarriage, stillbirth or ectopic pregnancy. Additionally, there are reports suggesting that pro-inflammatory cytokines are a risk factor for EP [52]. Further, we also observed that majority of the pathways (30 of 51) in the input gene set are classified as pathways for human diseases (Supplementary Table S5). All these observations signify that the immune-response related signaling pathways are strongly enriched in EP. Figure 4b shows the pathway similarity network for the KEGG pathways enriched in the input gene set (Methods). The pathway similarity network signifies the similarity of the gene sets between the KEGG pathways.

**Figure 4:**
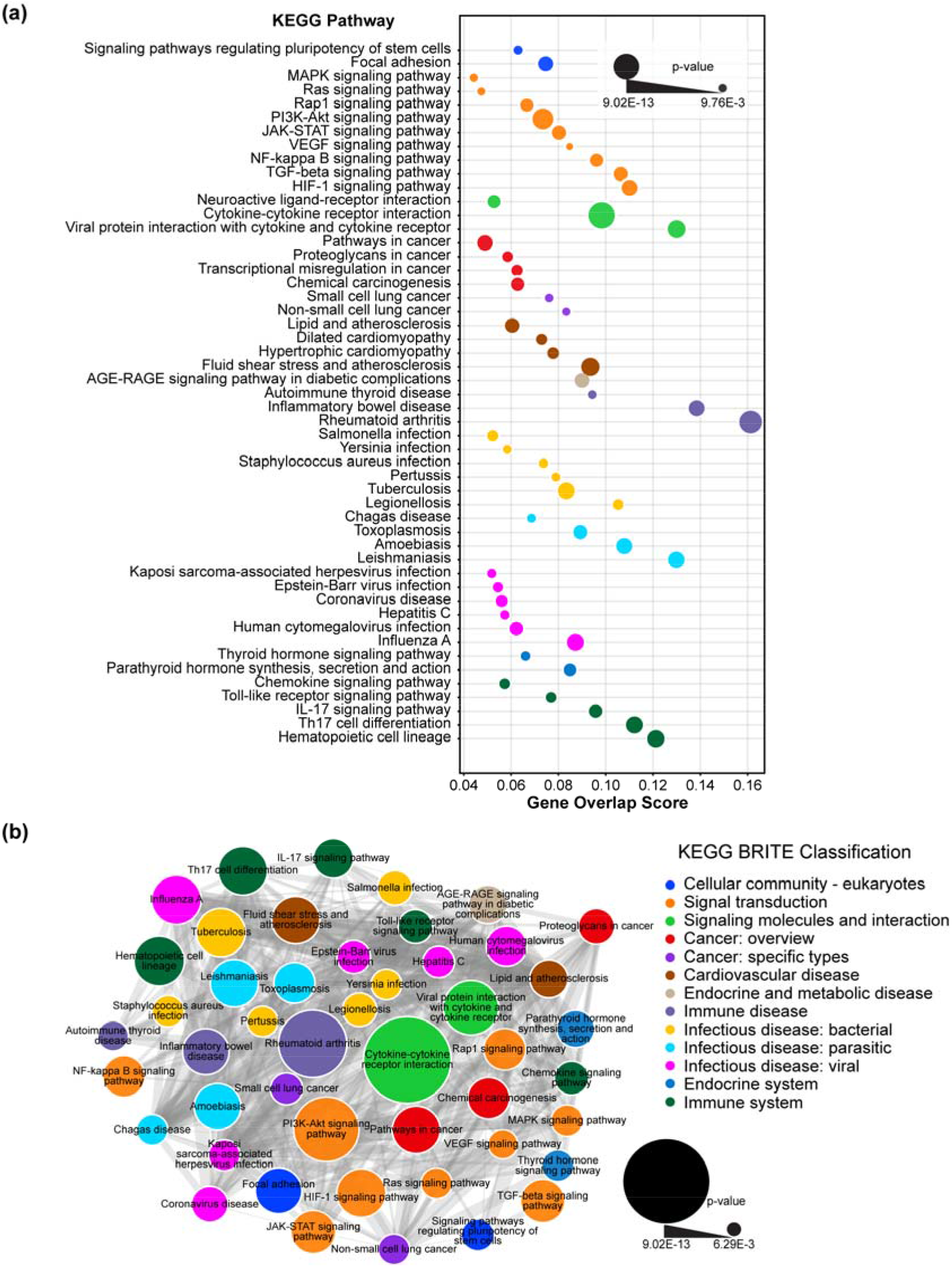
Pathway enrichment analysis to determine the pathways enriched in the 314 genes associated with EP (input gene set). (a) Bubble plot shows the 51 enriched KEGG pathways in the input gene set. The size of the bubble represents the p-value and position of the bubble is denoted by the overlapping genes with the input gene set. The bubbles are colored based on KEGG BRITE classification level 2. (b) Pathway similarity network of the KEGG pathways enriched in the input gene set. The similarity between two KEGG pathways is computed by the Jaccard index between the two sets of genes corresponding to both pathways. In the pathway similarity network, two pathways are connected if the similarity is > 0.1. The nodes in the pathway similarity network are colored based on the KEGG BRITE classification level 2.

### 3.4. PPI network highlights signaling related functional interactions

Protein-protein interaction (PPI) network is a general approach to reveal the functional interactions among a set of genes. We used the PPI network approach to reveal functional interactions among 314 genes associated with EP (Methods; Supplementary Table S6). Figure 5 shows the largest connected component of the first-order uterus-specific PPI network constructed starting with the 314 genes associated with EP. To better understand the interactions between the genes associated with EP, we visualized the zero-order uterus-specific PPI network within the first-order uterus-specific PPI network (Methods; Figure 5). In particular, we observed that the genes, FN1 and EGFR, have most connections in both the first-order and zero-order networks (Supplementary Tables S8 and S9). Moreover, we found that both FN1 and EGFR are involved in signaling processes. Therefore, the PPI network analysis, in conjunction with the enriched pathways, highlights the significance of signaling related interactions between genes associated with EP.

**Figure 5:**
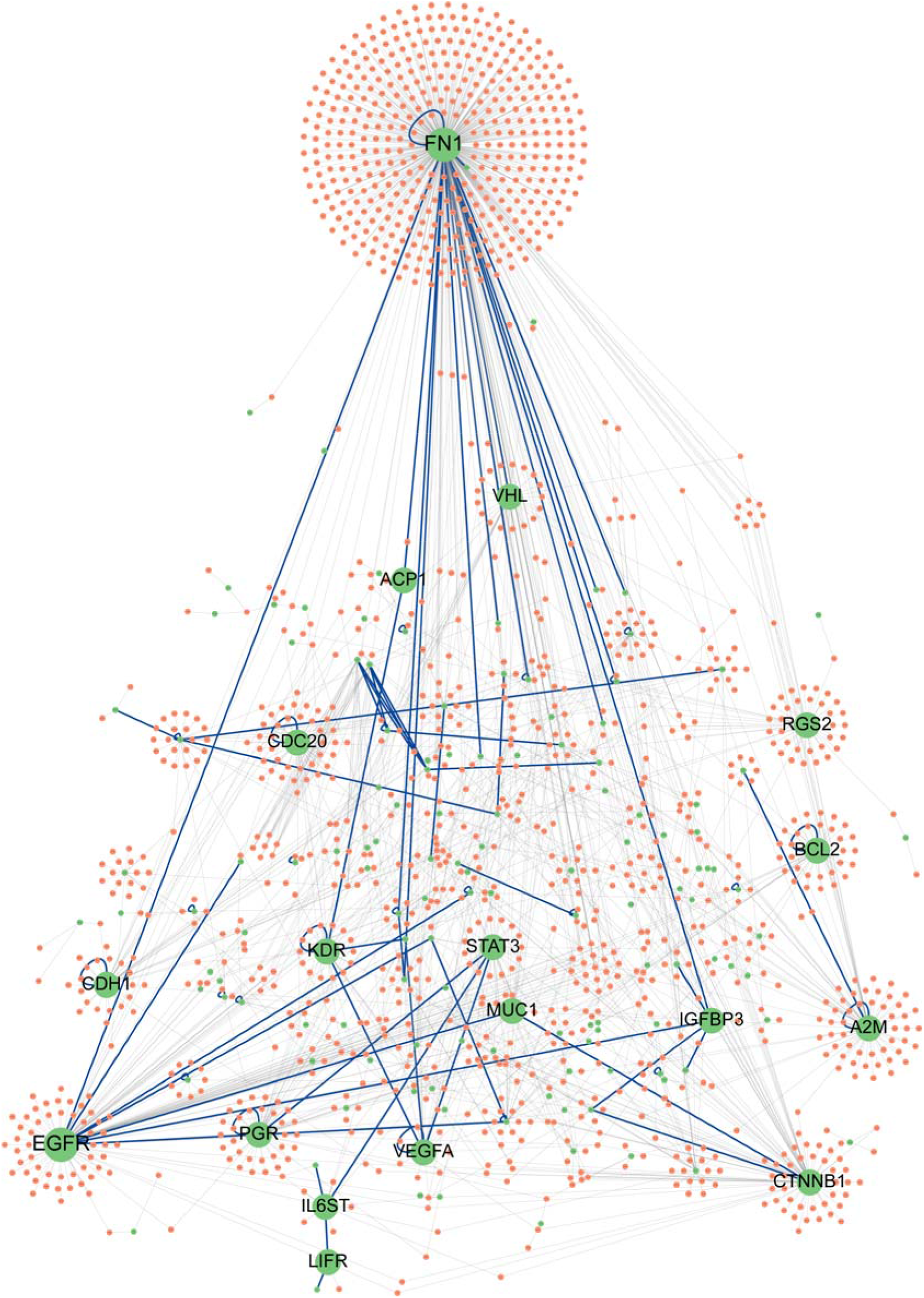
The largest connected component of the first-order uterus-specific PPI network constructed starting with the 314 genes associated with ectopic pregnancy (input gene set). In the PPI network, the nodes correspond to genes and the edges represent the interaction (undirected) between the connected genes. A node is colored in green if the corresponding gene belongs to the input gene set, and colored in brown otherwise. An edge is colored in blue if the two nodes connected by the edge belong to the input gene set (zero-order network), and colored in gray otherwise (first-order network). The top 10 nodes based on degree in both the first-order network and zero-order network are labelled with the corresponding NCBI gene symbols.

### 3.5. Ectopic pregnancy related gene markers and other maternal health disorders

We were interested to see the coverage of EPEK in comparison with other publicly available resources specific to EP. However, we did not find any specific resources on EP. So we relied on DisGeNET [35], one of the largest publicly available resource on various genedisease associations. We searched the DisGeNET database for EP-specific genes using the keywords ‘Ectopic Pregnancy’ and ‘Tubal Pregnancy’ and retrieved 50 genes that are reported to be associated with EP. We observed that there are only 9 overlaps with the gene set from EPEK.

During the literature survey on EP, we came across studies reporting genes that are also associated with other maternal health disorders [53–55]. Therefore, we were motivated to check for resources specific to other maternal health disorders that might have captured the genes associated with EP. We identified curated resources like PrecocityDB [10] associated with precocious puberty, PCOSKB [11] associated with polycystic ovary syndrome, Endometriosis Knowledgebase [12] associated with endometriosis, and CCDB [13] associated with cervical cancer. Further, Barbitoff *et al*. [56] have manually curated the list of genes associated with 4 common pregnancy complications namely, preeclampsia, gestational diabetes, placental abruption and preterm birth from public resources. Importantly, we observed that there are few expression-centric resources available for maternal health diseases in the public domain.

Thereafter, we collected the genes associated with these 8 maternal health disorders from the above-mentioned resources and analyzed the overlaps with the genes associated with EP (Figure 6). From Figure 6, we observed that EPEK captures 194 genes (~62% of input gene set) which are unique, i.e. there is no overlap of these 194 genes with the gene sets of the other 8 maternal health disorders captured in the above-mentioned resources. In sum, this analysis suggests that the majority of the differentially expressed genes in EP that are compiled in EPEK are unique, and are not implicated in public databases with many of the maternal health disorders analyzed here.

**Figure 6:**
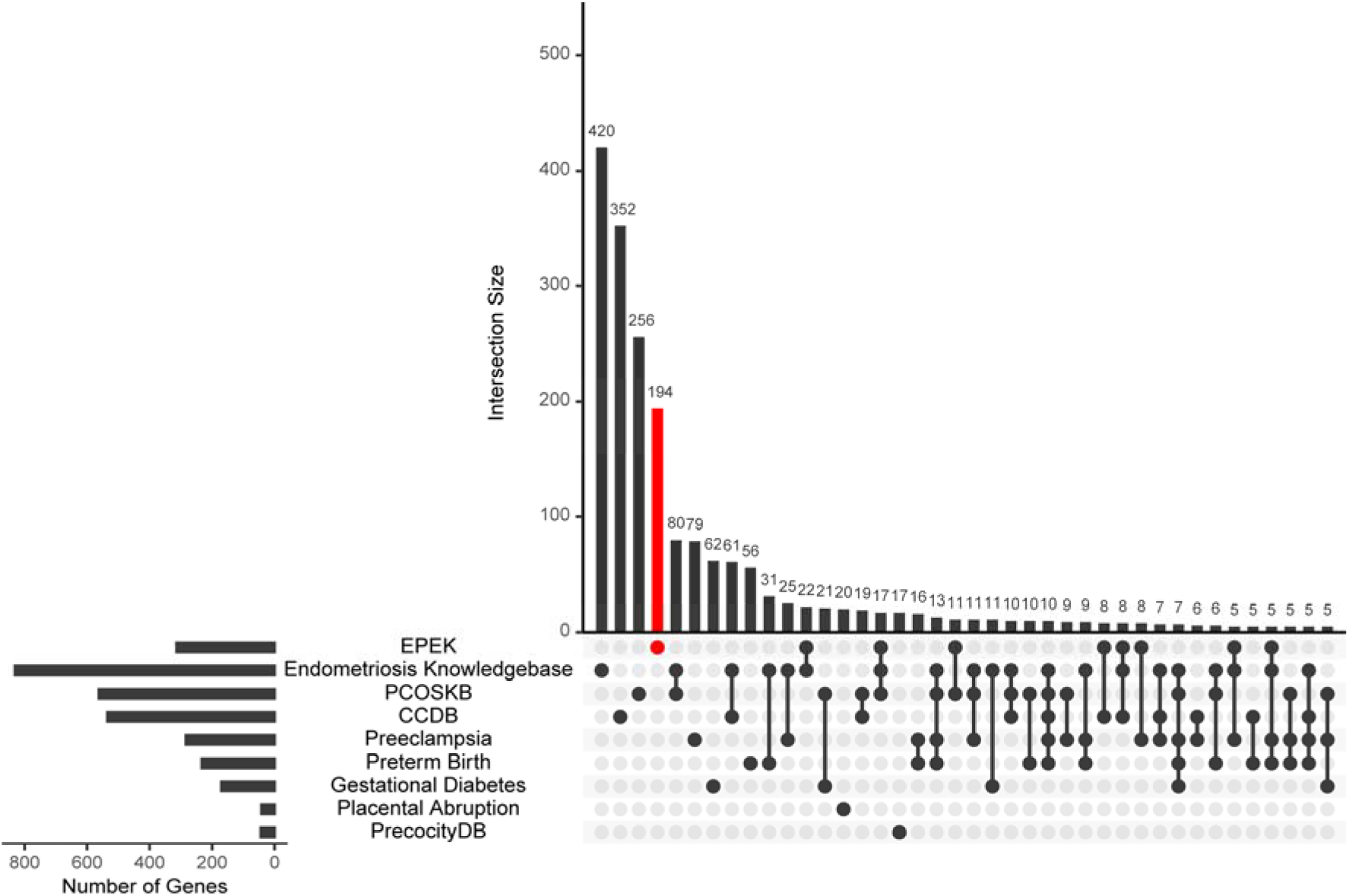
UpSet [64] plot illustrating the overlaps among genes compiled in 9 maternal health resources considered here. The number of genes specific to each intersection is provided on the top of the vertical bar in the plot. The vertical bar colored in red corresponds to the unique number of genes compiled in EPEK. Further, the horizontal bar corresponds to the number of genes present in each resource.

From the above analysis, a question may arise on the need for disease-specific databases when a resource like DisGeNET is already available. It is important to note that DisGeNET compiles gene-disease associations from various sources and is largely dependent upon automated text mining. In contrast, resources like EPEK and other disease-specific resources mentioned above are manually curated. In our effort to build EPEK, we realized that manual curation helps in managing the under-represented or over-represented data, and in filtering out spurious data. Further, in the comparison of EPEK with DisGeNET, manual curation has led to compilation of more than six times of the genes covered in DisGeNET and gives a more comprehensive dataset to date on EP for academic research.

### 3.6 Disease enrichment suggests inflammation as a probable risk factor for EP

Disease enrichment analysis helps in finding the probable comorbidity risk. In this study, we performed the disease enrichment analysis to find diseases enriched in the 314 genes associated with EP (Methods; Supplementary Table S7). Figure 7a shows the bubble plot of the 49 enriched diseases in the input gene set. In Figure 7a, we identified ‘Inflammation’ to be strongly enriched while also having the largest overlap score. Inflammation is the biological response of the tissue to different harmful stimuli. It has been reported in literature that mothers infected with pelvic inflammatory disorder have a higher risk of tubal ectopic pregnancy over healthy mothers [57–61]. Moreover, we observed that a large number of enriched diseases in the input gene set are categorized as neoplasms (Figure 7a). Neoplasm progression is attributed to the chronic inflammation of the tissues [62], thereby suggesting that inflammation could be a risk factor for EP. Figure 7b shows the disease-disease similarity network of the diseases from DisGeNET enriched in the input gene set (Methods). The disease-disease similarity network signifies the similarity of the gene sets associated with the diseases.

**Figure 7:**
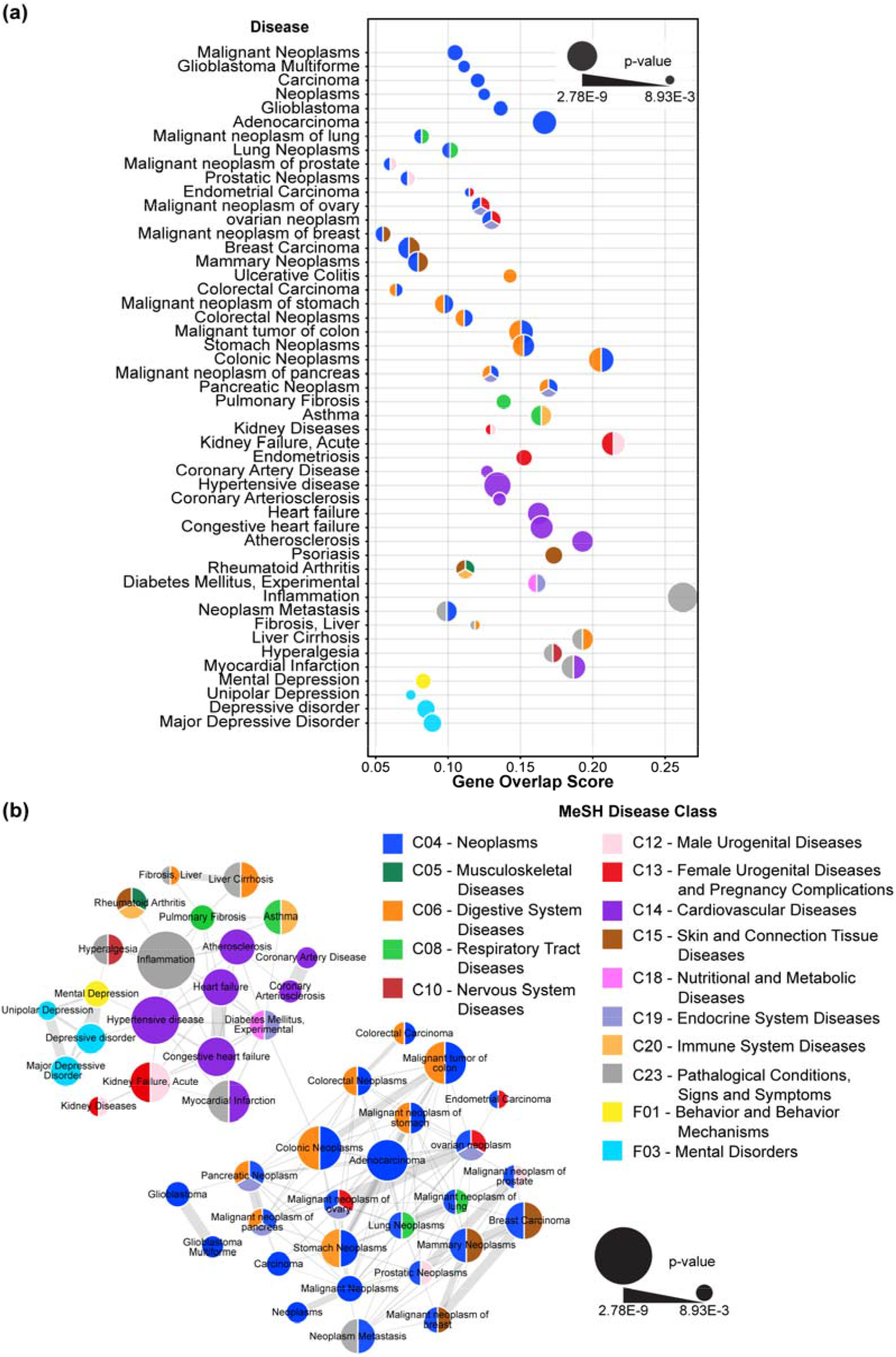
Disease enrichment analysis to determine the diseases enriched in the 314 genes associated with EP (input gene set). (a) Bubble plot shows the 49 diseases from DisGeNET enriched in the input gene set. The size of the bubble represents the p-value, and the position gives the overlap with the input gene set. The bubbles are colored based on disease classification from MeSH. (b) Disease-disease similarity network of the enriched diseases in the input gene set. The similarity between any two diseases is computed by the Jaccard index between the two sets of genes corresponding to both diseases. In the disease-disease similarity network, two diseases are connected if the similarity is > 0.1. The nodes in the disease-disease similarity network are colored based on the disease classification from MeSH.

### 4. Discussions and Conclusions

The present study reports the first dedicated resource on the expression profile of ectopic pregnancy in humans, namely EPEK. EPEK is a manually curated resource that involved an extensive and systematic manual effort in compiling the experimental information on 314 differentially expressed genes, 17 metabolites and 3 SNPs associated with EP from published articles. EPEK also includes the various annotations for its records that are collected from standard databases. The enrichment analyses on the genes compiled in EPEK showed a strong involvement of genes in different cell signaling mechanisms. The PPI network also highlighted that the interactions are signaling related, suggesting that abnormal cellular signaling is a prominent phenotype associated with EP.

Our analysis on the gene set in EPEK also highlighted the clinical potential of genetic markers involved in cellular signaling. Noting that exosomes are known to carry signaling molecules and are clinically relevant, we identified three miRNAs and a protein coding gene by comparing EPEK with existing datasets on exosomes secreted from the uterus. The limitation of this comparative analysis is that the existing exosome datasets are not specific to EP. Till date, there is no comprehensive study on exosomes specific to EP, and our study motivates the need for identification of EP-specific exosome markers to facilitate early diagnosis of ectopic pregnancy.

Computational resources specific to maternal health disorders have been developed to assist the ongoing efforts in elucidating the underlying mechanisms of the disorders. A comparative analysis of EPEK with these different resources not only revealed that the different maternal health disorders have a substantial set of unique genes, but also showed the potential of EPEK to explain the EP-specific mechanisms. In an earlier attempt to understand the genes associated with EP, Liu and Zhao [63] had compiled a set of 269 genes based on automated text mining from published articles. While the gene set compiled by Liu and Zhao did not contain any miRNA, it did not show a substantial overlap with the gene set in EPEK (only 69 overlapping genes). In contrast, EPEK not only compiles 314 genes, but also includes metabolites and SNPs that have been manually curated from published articles. This underscores the importance of EPEK as a dedicated resource for ectopic pregnancy in humans.

In conclusion, EPEK has the potential to help in the design of models that can predict the likelihood of the development of ectopic pregnancy. EPEK can also aid in data-driven hypothesis generation for studies focussed on identification of novel therapeutic targets in the treatment of ectopic pregnancy.

## Data and code availability

EPEK database is accessible via the associated website: https://cb.imsc.res.in/epek. The compiled information in EPEK is made available under a Creative Commons Attribution-NonCommercial 4.0 (CC BY-NC 4.0) International License (http://creativecommons.org/licenses/by-nc/4.0/). The computer codes used to perform enrichment analysis and to generate associated plots are available via the associated GitHub repository: https://github.com/asamallab/EPEK.

## Acknowledgements

We would like to thank Kishan Kumar for help with figures and Anshu Bhardwaj for insightful discussions. A.S. would like to acknowledge funding from the Department of Atomic Energy (DAE), Government of India and the Max Planck Society, Germany via a Max Planck Partner Group in Mathematical Biology. The funders have no role in study design, data collection, data analysis, manuscript preparation or decision to publish.

## CRediT author contribution statement

**Ananya Natarajan:** Conceptualization, Data curation, Formal analysis, Writing. **Nikhil Chivukula:** Conceptualization, Data curation, Formal analysis, Software, Visualization, Writing. **Gokul Balaji Dhanakoti:** Data curation, Formal analysis, Software, Visualization, Writing. **Ajaya Kumar Sahoo:** Formal analysis, Writing. **Janani Ravichandran:** Conceptualization, Data curation, Supervision, Formal analysis, Writing. **Areejit Samal:** Conceptualization, Supervision, Formal analysis, Writing.

## Supplementary Tables

**Table S1:** List of 314 genes identified from published articles, associated with ectopic pregnancy. For each gene, we provide the gene identifier and the gene symbol from NCBI (https://www.ncbi.nlm.nih.gov/gene/). Further, we provide the expression type and the PubMed identifiers (separated by ‘|’ symbol) of the published articles.

**Table S2:** List of 17 metabolites identified from published articles, associated with ectopic pregnancy. For each metabolite we provide the CAS identifier from ChemIDplus (https://chem.nlm.nih.gov/chemidplus/) database, metabolite name, and PubMed identifiers (separated by ‘|’ symbol) for published articles.

**Table S3:** List of 3 SNPs identified from published articles, associated with ectopic pregnancy. For each SNP, we provide the SNP identifier from the dbSNP (https://www.ncbi.nlm.nih.gov/snp/) database, gene symbol and gene identifier from NCBI (https://www.ncbi.nlm.nih.gov/gene/), and PubMed identifiers for published articles.

**Table S4:** List of 30 GO terms enriched in the 314 genes associated with ectopic pregnancy (input gene set). The 30 GO terms are categorized into 3 ontologies namely, Biological Process, Molecular Function and Cellular Component. For each GO term, we provide the GO term identifier, GO term name, ontology, p-value, number of genes from the input gene set associated with the GO term, and NCBI gene symbols of the associated genes (separated by ‘|’ symbol) from GOnet (https://tools.dice-database.org/GOnet/) web-application.

**Table S5:** List of 51 KEGG pathways enriched in the 314 genes associated with ectopic pregnancy (input gene set) using Enrichr (https://maayanlab.cloud/Enrichr/). For each KEGG pathway, we provide KEGG BRITE classification (Level 1 and Level 2), number of genes associated with KEGG pathway, number of genes in KEGG pathway overlapping with the input gene set, overlap score, p-value and NCBI gene symbol of overlapping genes (separated by ‘|’ symbol).

**Table S6:** The edge file for the largest connected uterus-specific PPI netowrk derived from NetworkAnanlyst 3.0 (https://www.networkanalyst.ca/NetworkAnalyst/home.xhtml). For each edge, we provide the NCBI gene identifiers and the NCBI gene symbols for the two nodes (Node 1 and Node 2) connected by the edge. An edge is considered as an edge in zero-order network if the two nodes connected by the edge belong to the 314 genes associated with ectopic pregnancy. For each edge, we assigned a value of 1 if it belongs to the zero-order network, 0 otherwise.

**Table S7:** List of 49 enriched diseases in the 314 genes associated with ectopic pregnancy (input gene set) from DisGeNET (https://www.disgenet.org/) database. For each disease, we provide the disease code, disease name, disease classes (seperated by ‘|’ symbol), number of genes associated with the disease, number of genes overlapping with the input gene set, overlap score, p-value, and NCBI gene symbol of overlapping genes (separated by ‘|’ symbol).

**Table S8:** The top 10 genes ordered based on the degree in the first-order uterus-specific PPI network of the 314 genes associated with ectopic pregnancy. For each gene, we provide the NCBI gene identifier, NCBI gene symbol, the degree in first-order PPI network, presence in resources specific to other maternal health disorders (seperated by ‘|’ symbol), and the signaling pathways identified form KEGG database (https://www.genome.jp/kegg/) (seperated by ‘|’ symbol).

**Table S9:** The top 10 genes ordered based on the degree in the zero-order uterus-specific PPI network of the 314 genes associated with ectopic pregnancy. For each gene, we provide the NCBI gene identifier, NCBI gene symbol, the degree in zero-order PPI network, presence in resources specific to other maternal health disorders (seperated by ‘|’ symbol), and the signaling pathways identified form KEGG database (https://www.genome.jp/kegg/) (seperated by ‘|’ symbol).

